# Coordination of gene expression noise with cell size: analytical results for agent-based models of growing cell populations

**DOI:** 10.1101/2020.10.23.352856

**Authors:** Philipp Thomas, Vahid Shahrezaei

## Abstract

The chemical master equation and the Gillespie algorithm are widely used to model the reaction kinetics inside living cells. It is thereby assumed that cell growth and division can be modelled through effective dilution reactions and extrinsic noise sources. We here re-examine these paradigms through developing an analytical agent-based framework of growing and dividing cells accompanied by an exact simulation algorithm, which allows us to quantify the dynamics of virtually any intracellular reaction network affected by stochastic cell size control and division noise. We find that the solution of the chemical master equation – including static extrinsic noise – exactly agrees with the agent-based formulation when the network under study exhibits *stochastic concentration homeostasis*, a novel condition that generalises concentration homeostasis in deterministic systems to higher order moments and distributions. We illustrate stochastic concentration homeostasis for a range of common gene expression networks. When this condition is not met, we demonstrate by extending the linear noise approximation to agent-based models that the dependence of gene expression noise on cell size can qualitatively deviate from the chemical master equation. Surprisingly, the total noise of the agent-based approach can still be well approximated by extrinsic noise models.

## I. INTRODUCTION

Cells must continuously synthesise molecules to grow and divide. At a single cell level, gene expression and cell size are coordinated but heterogeneous which can drive phenotypic variability and decision making in cell populations^1–5^. The interplay between these sources of cell-to-cell variability is not well understood since they have traditionally been studied separately. A general stochastic theory integrating size-dependent biochemical reactions with the dynamics of growing and dividing cells is hence still missing.

Many models of noisy gene expression and its regulation are based on the chemical master equation that describes the stochastic dynamics of biochemical reactions in a fixed reaction volume^6–8^. The small scale of compartmental sizes of cells implies that only a small number of molecules is present at any time leading to large variability of reaction rates from cell to cell, commonly referred to as gene expression noise^9–11^. Another factor contributing to gene expression noise is the fact that cells are continuously growing and dividing causing molecule numbers to (approximately) double over the course of a growth-division cycle. A common approach to account for cell growth is to include extra degradation reactions that describe dilution of gene expression levels due to cell growth^9–13^ akin to what is done in deterministic rate equation models^14,15^. We will refer to this approach as the *effective dilution model* (EDM, see Fig. 1a). However, little is known of how well this approach represents the dependence of gene expression noise on cell size observed in a growing population.

**FIG. 1.**
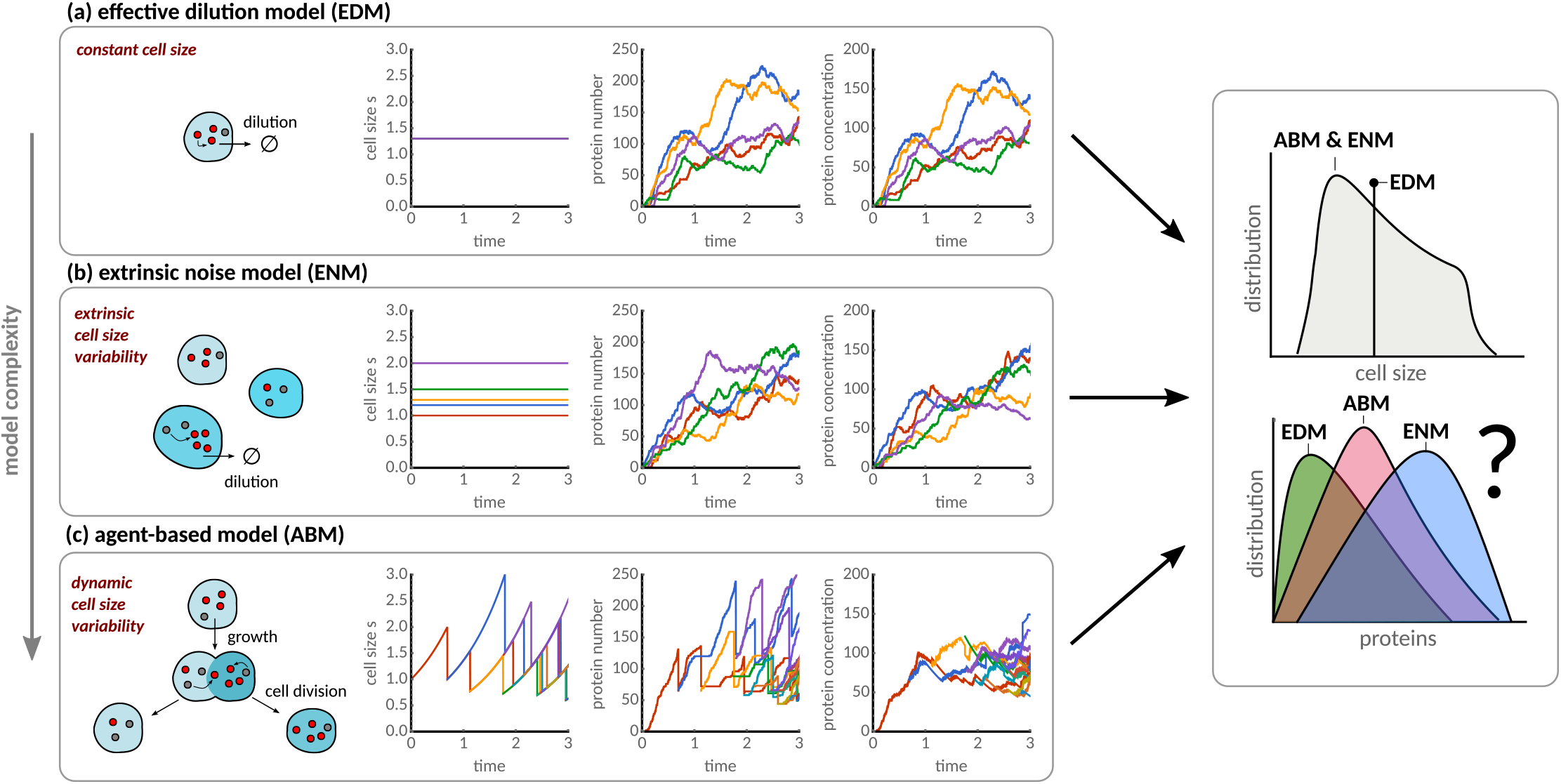
Modelling approaches for cell size dependence of gene expression. **(a)** The *effective dilution model* describes cells at constant size with intracellular reactions coupled to effective dilution reactions. **(b)** The *extrinsic noise model* incorporates static cell size variability as a source of extrinsic noise coupled with effective dilution models **(c)** The *agent-based approach* models intracellular reactions occurring across a growing and dividing cell population without the need for effective dilution reactions.

Cells achieve concentration homeostasis through coupling reaction rates to cell size via highly abundant upstream factors like cell cycle regulators, polymerases or ribosomes that approximately double over the division cycle^3,16,17^. Cell size fluctuates in single cells, however, providing a source of extrinsic noise in reaction rates that can be identified via noise decompositions^18,19^. A few studies combined EDMs with static cell size variations as an explanatory source of extrinsic noise^20–22^. In brief, the total noise in these models amounts to intrinsic fluctuations due to gene expression and dilution, and extrinsic variation across cell sizes in the population. We refer to this class of models as *extrinsic noise models* (ENMs, see Fig. 1b). Yet it remains unclear how reliably these effective models describe cells that continuously synthesise molecules, grow and divide.

An increasing number of studies are investing efforts towards quantifying the dependence of gene expression noise on cell cycle progression and growth, either experimentally via ergodic principles or pseudo-time^23,24^ and time-lapse imaging^22,25,26^ or theoretically through noise decomposition^27–29^, master equations including cell cycle dynamics^4,17,30–35^ and agent-based approaches including age-structure of growing populations^35–40^. The essence of *agent-based models* (ABMs) is that each cell in a population is represented by an agent whose physiological state is tracked along with their molecular reaction networks. In principle, these models are able to predict gene expression distributions of cells progressing through well-defined cell cycle states as measured by time-lapse microscopy and snapshots of heterogeneous populations. The unprecedented detail of these models must cast doubt on the predictions of master equation models (EDMs and ENMs) in which growth and division are modelled by effective dilution reactions. Yet it is presently unclear why these effective models have fared reasonably well in predicting gene expression noise reported by single-cell experiments^10,17,41^.

Nevertheless, most ABMs still ignore cell size, a major physiological factor affecting both intracellular reactions and cell division dynamics alike. Since cell size varies at least two-fold as required by size homeostasis in a growing population, and it scales some reaction rates as required by concentration homeostasis, it is expected that cell size must significantly contribute to gene expression variation across a population. In this article, we bridge the gap between the chemical master equation and agent-based approaches by integrating cell size dynamics with the stochastic kinetics of molecular reaction networks.

The outline of the paper is as follows. First, we explain the analytical framework for EDMs, ENMs and ABMs (II). Then we introduce the concept of *stochastic concentration homeostasis*, a rigorous condition under which the chemical master equations of the EDM and ENM agree exactly with the ABM (Sec. III A). This new condition is met by some but not all common models of gene expression. We show that when these conditions are not met, the effective models agree with the ABM only on average (Sec. IIIB). To address this problem, we propose a comprehensive theoretical framework extending the linear noise approximation to agent-based dynamics with which we quantify cell size scaling of gene expression in growing cells (Sec. III C). Our findings indicate that the EDM can qualitatively fail to predict this dependence but our novel approximation method accurately describes gene expression noise in the presence of cell size control variations and division errors. We further show that ENMs present surprisingly accurate approximations for the total noise statistics (Sec. III D).

## II. METHODS

We consider a biochemical reaction network of *N* molecular species *S* = (*S*_1_, *S*_2_,…, *S_N_*)^*T*^ embedded in a cell of size *s*. The network then has the general form:

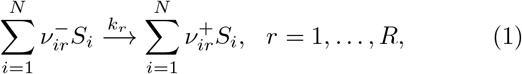

where 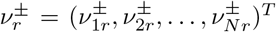 are the stoichiometric coefficients and *k_r_* is the reaction rate constant of the *r^th^* reaction. In the following, we outline deterministic, effective dilution and extrinsic noise models and develop a new agent-based approach coupling stochastic reaction dynamics to cell size in growing and dividing cells (Fig. 1).

### A. Effective dilution models, extrinsic noise models and the chemical master equation

#### 1. Rate equation models and concentration homeostasis

Deterministically, the vector of molecular concentrations 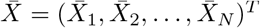 is governed by rate equation models in balanced growth conditions. The balanced growth condition states that there exists a steady state between reaction and dilution rates

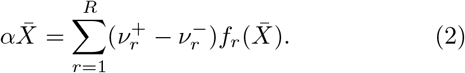

Here, 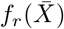 are macroscopic reaction-rate functions and *α* is the exponential growth rate of cells determining the dilution rate due to growth. Since these quantities are independent of cell size, the balanced growth condition (2) implies concentration homeostasis in rate equation models.

#### 2. Effective dilution model

The chemical master equation^6^ and equivalently the stochastic simulation algorithm^7^ are state-of-the-art stochastic models of reaction kinetics inside cells. Although well-established, they are strictly valid only when describing cellular fluctuations at constant cell size *s*. A straight-forward approach to circumvent this limitation is to supplement (1) by additional degradation reactions of rate α that model dilution of molecules due to cell growth:

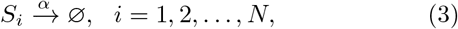

akin to what is traditionally for reaction rate equations (2). The chemical master equation of this *effective dilution model* (EDM) then takes the familiar form

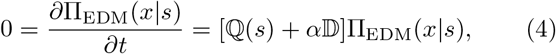

governing the conditional probability of molecule numbers *x* = (*x*_1_, *x*_2_,…, *x_N_*)^T^ of the species *S* in a cell of size s and where

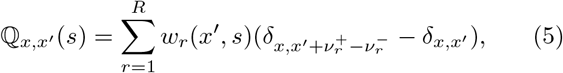

are the elements of the transition matrix of the molecular reactions (1) and we included the extra dilution reactions (3) via 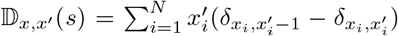. We are here interested in the stationary solution and hence set the time-derivative in Eq. (4) to zero. Such effective models are motivated through the fact^6,42^ that when the microscopic propensities *w_r_* are linked to the macroscopic rate functions *f_r_* of the rate equation models via mass-action kinetics

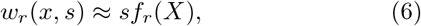

where *X* = *x/s* is the concentration, the mean concentrations of EDMs follow the concentrations *X* of the rate equations (2) (see Sec. IIA 4).

#### 3. Extrinsic noise model

A common way to incorporate static size variability between cells in the model is to consider cell size *s* to be distributed across cells according to a cell size distribution Π(*s*). We will refer to this approach as the *extrinsic noise model* (ENM), which leads to a mixture model of concentrations *X* = *x/s*,

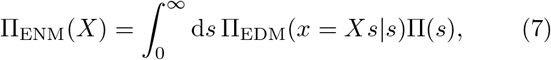

and analogous expressions for the molecule number distributions.

#### 4. Analytical solutions and noise decomposition

The advantage of the EDM and ENM is that its noise statistics can be approximated in closed-form using the linear noise approximation^6,43,44^. In this approximation, the mean concentrations are approximated by the solution 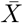 of the rate equations (2) and the probability distribution Π_EDM_(*x*|*s*) is approximated by a Gaussian. In the same limit, the covariance matrix Σ_*Y*_ can be decomposed into intrinsic and extrinsic components, 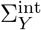 and 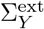, using the law of total variance^18,19^

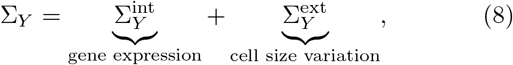

which correspond to molecular fluctuations due to gene expression and cell size variation, respectively, for *Y* ∈ {EDM, ENM}. Specifically, for molecule numbers *x*, we have 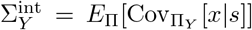 and 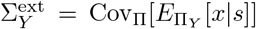, where *E*_Π_ denotes the expectation value with respect to the distribution Π, and analogously for concentrations. The intrinsic components 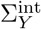 satisfy a Lyapunov equation called the linear noise approximation:

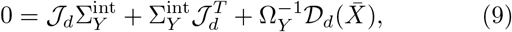

where Ω_*Y*_ has to be chosen depending on whether concentration or number covariances are of interest:

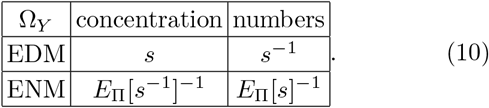

The matrix 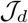 is the Jacobian of the rate equations (2) and 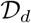 denotes the diffusion matrix obeying

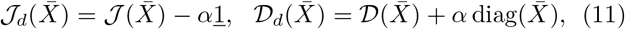

where 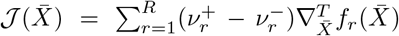 and 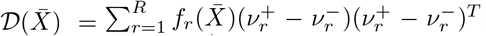. The extrinsic components 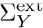 follow from the dependence of the mean on cell size, which features only in the molecule number variance of the ENM:

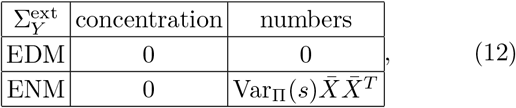

where the last cell follows from 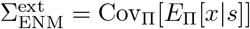 with 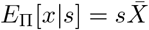.

As a concrete example, we consider transcription of mRNAs with a size-dependent transcription rate that are translated into stable proteins:

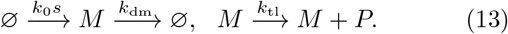

We then account for dilution through the additional reactions

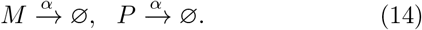

The mean protein concentration is given by 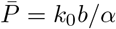 and the coefficient of variation predicted by the EDM and ENM models follow the familiar expression^10^

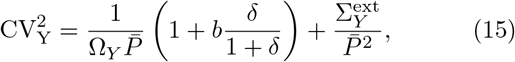

where we account for size-variability via Ω_*Y*_ and 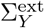 given by Eqs. (10) and (12), respectively, and the parameters

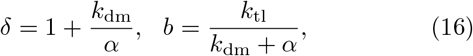

correspond to the ratio of mRNA and protein degradation/dilution rates and the translational burst size, respectively. From Eq. (15), (10) and (12), it is clear that size variation acts on the intrinsic noise component of molecule concentrations (via 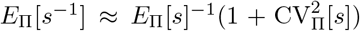) but the extrinsic noise component of molecule numbers (via 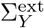 (12)).

### B. Agent-based modelling

Little is known about the accuracy of EDMs and ENMs in predicting cellular noise in growing populations. In the following, we introduce an agent-based modelling approach that serves as a gold standard to assess the validity of these effective models. The ABM represents cells as agents that progressively synthesise molecules via intracellular reactions (1), grow in size and undergo cell division. Every division gives rise to two daughter cells of varying birth sizes, each of which inherits a proportion of molecules from the mother cell via stochastic size-dependent partitioning at division.

The ABM simulation algorithm is given in Box 1, which combines the First-Division algorithm, previously introduced for agent-based cell populations^38^, with the Extrande method adapted to simulate reaction networks embedded in a growing cell^45^. In the following, we describe the exact analytical framework with which we characterise the snapshot distributions that underlie such a population of agents.

#### Box 1: First-Division Algorithm for agent-based simulations of size-dependent gene regulatory networks

Exact simulation algorithm of general stochastic reaction networks within growing cells (agents) undergoing binary cell division according to cell size control rules^46,50,52^. The algorithm combines the Extrande method^45^ for simulating reaction networks embedded in a growing cell and the First-Division algorithm^38^ for the population dynamics. The state of each cell is given by birth time *t*_0_, birth size *s*_0_, present cell size *s* and the vector of molecule numbers *x*.

##### Algorithm 1

First-division algorithm simulating agent-based population dynamics

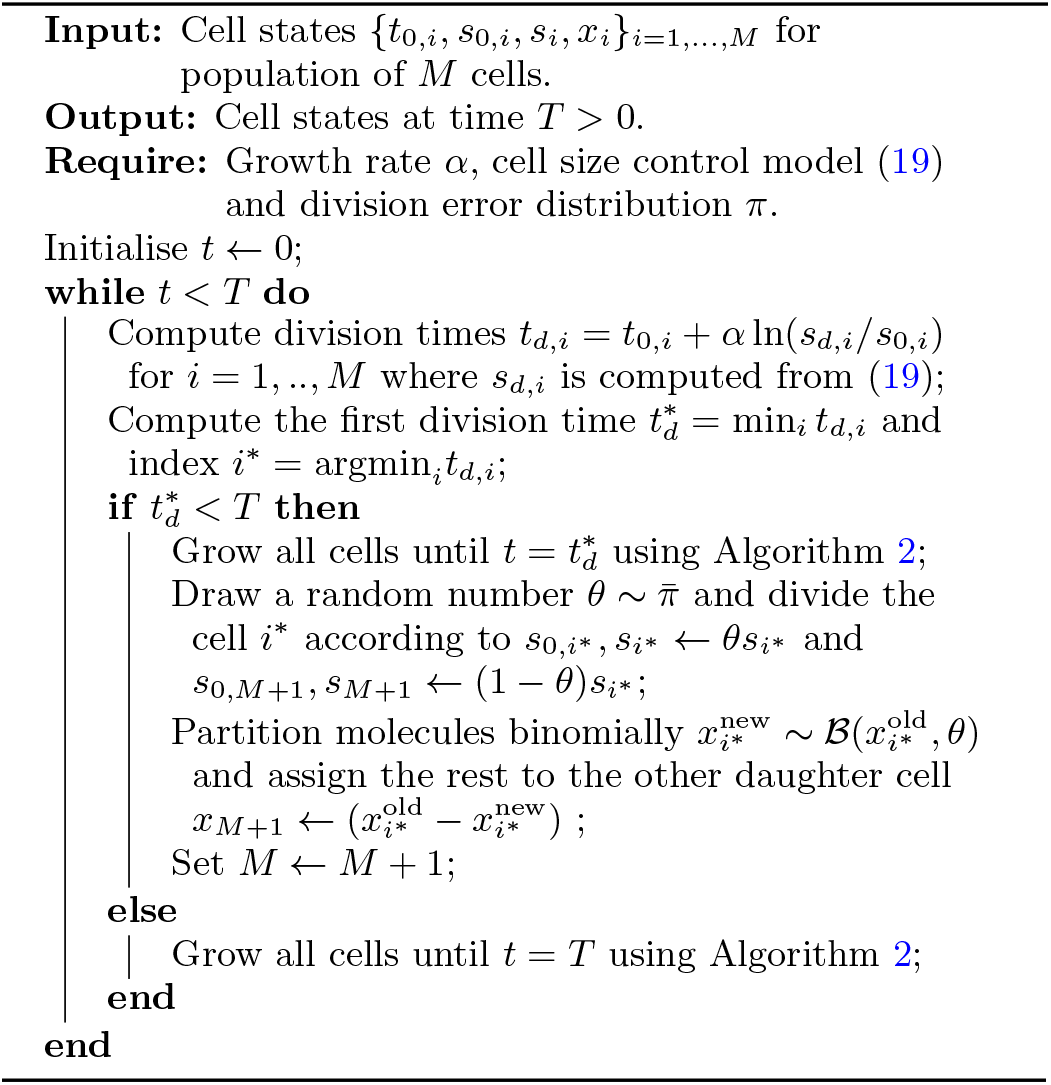

##### Algorithm 2

Extrande algorithm simulating reaction networks in a growing cell

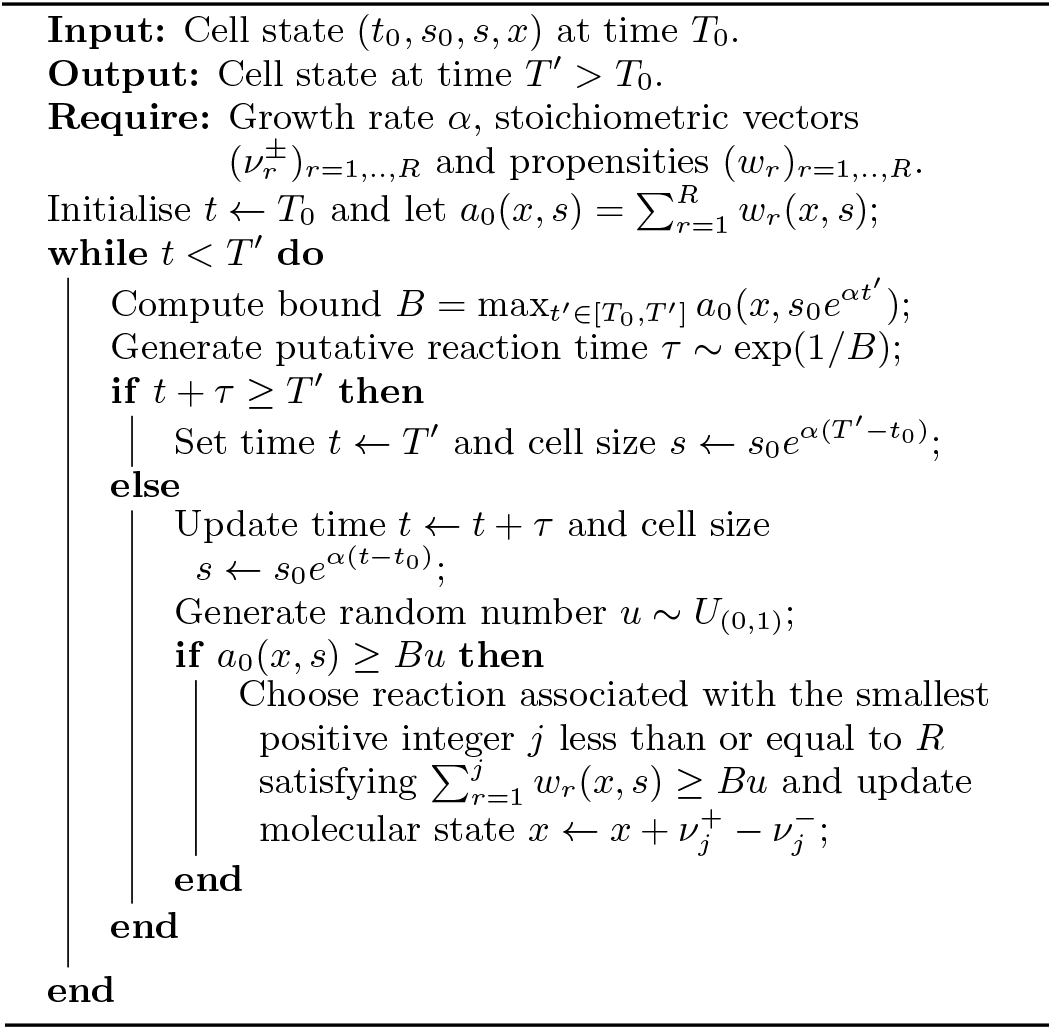

#### Master equation for agent-based populations

We consider the number of cells *n*(*τ,s,x,t*) with age *τ* (time since the last division), cell size s and molecule counts *x* in a snapshot at time *t*, which evolves as

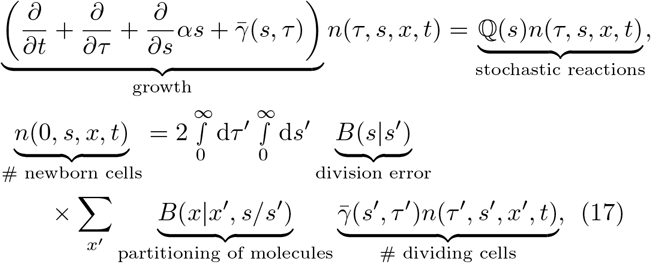

and describes cell growth, stochastic reaction kinetics and a boundary condition for cell division that ensures that the number of newborn cells is twice the number of dividing cells after partitioning their size and molecular contents. These evolution equations have been derived in^38,39^ for age-dependent snapshots but here we extend such agent-based models to include also cell size dynamics and size-dependent reaction dynamics. We allow for the following generalisations: (i) size increases exponentially in single cells, (ii) cells divide with rate 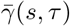 that is both size- and age-dependent, (iii) the transition matrix 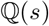 of the molecular reactions depends on cell size s via the propensities (see definition after Eq. (4)), and (iv) the molecular partitioning kernel *B*(*x*|*x′, s/s′*) depends on the inherited size fraction *s/s′* of a daughter cell. We now describe in detail how we model the individual noise sources associated with cell size control, division errors, and molecule partitioning.

##### a. Cell size control fluctuations

Recent studies^46,47^ have shown that the distribution of sizes with which cells divide does not explicitly depend on cell age but on the birth size *s*_0_. Assuming that 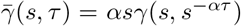, where *γ*(*s, s*_0_) is the division rate per unit size (see also^48,49^), the division-size distribution is given by

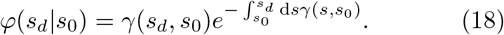

As a concrete example of (18) we consider a model where the division size is linearly related to birth size^49–51^

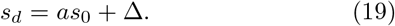

The division rate can be calculated from the distribution 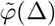 of the noise term Δ in (19) via 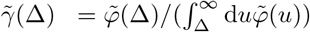 and setting 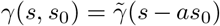, which gives the correct division-size distribution 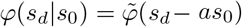 as expected. The model generalises the sizer (*a* = 0) to concerted cell size controls such as the adder (*a* = 1) and timer-like (2 > *a* > 1) models^46,47,52^. In the following, we will refer to CV_*φ*_[Δ] as the size-control noise.

##### b. Division errors

After division, size is partitioned between cells and the birth size of the two daughter cells is obtained from 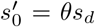 and 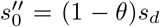 where *θ* is the inherited size fraction, a random variable between 0 and 1 with distribution 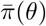 (see Box 1). This can be modelled using the division kernel

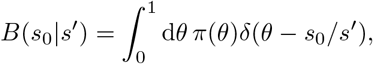

where 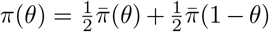 including the case of asymmetric division. We will refer to CV_*π*_[*θ*] as the division error about the centre 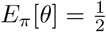.

##### c. Molecule partitioning at cell division

The partitioning kernel *B*(*x*|*x′, θ*) denotes the probability that a cell inherits x molecules from a total of *x′* molecules from its mother and this probability depends on the daughter’s inherited size fraction *θ*. We assume that cells are sufficiently well mixed and each molecule is partitioned independently with probability *θ* such that the division kernel is binomial

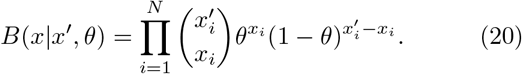

To make analytical progress, we assume that the population establishes a long-term stationary distribution Π(*s, s*_0_, *x*) characterising the fraction of cells with molecule numbers *x*, cell size s and birth size *s*_0_ that is invariant in time. To this end, we let *n*(*τ, s, x, t*) ∝ *e^αt^*Π(*s, τ, x*) and change variables from cell age *τ* to birth size *s*_0_ such that Π(*s, s*_0_, *x*) = (*αs*)^-1^Π(*s, τ* = ln(*s/s*_0_)/*α, x*). We find that this transformation reduces the PDE (17) to an integro-ODE:

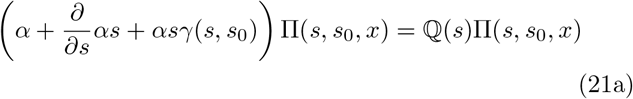

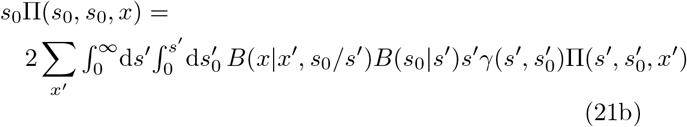

We finally characterise the marginal cell size distribution Π(*s, s*_0_) and the conditional molecule number distribution Π(*x*|*s, s*_0_) via Bayes’ formula

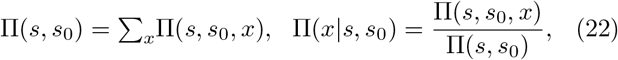

which together provide the full information about the population snapshot.

#### Cell size distribution

The evolution of the size distribution Π(*s, s*_0_) is obtained by summing Eqs. (21) over all possible *x*, which yields:

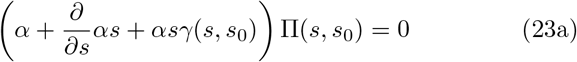

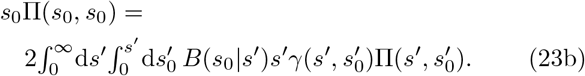

Eqs. (23) can be solved analytically

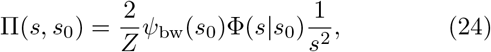

where *ψ*_bw_(*s*_0_) is the birth size distribution in a backward lineage (see^48^ for details), 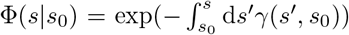 is the probability that a cell born at size *s*_0_ has not divided before reaching size *s*, and 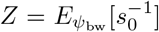 is a normalising constant.

#### Molecule number distributions for cells of a certain size

The conditional molecule number distribution Π(*x|s, s*_0_) gives the probability to find the molecule numbers *x* in a cell of size s that was born at size *s*_0_ and satisfies

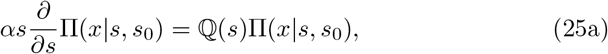

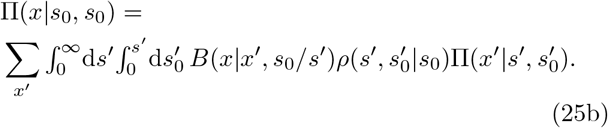

Eqs. (25) follow directly from substituting Eq. (22) into (21) and using (18) and (23). The solution of these equations depends implicitly on the ancestral cell size distribution *ρ*,

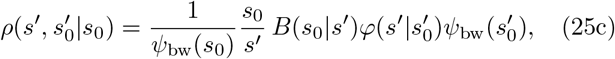

that gives the probability of a cell born at size *s*_0_ having an ancestor with division size *s′* and birth size 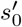. The main difference between the molecule number distributions of the ABM and the EDM/ENM is the boundary condition at cell division, which as we shall see can have a significant effect on the reaction dynamics.

## III. RESULTS

We here introduce the concept of *stochastic concentration homeostasis* (SCH) as a generalisation of concentration homeostasis in deterministic systems (see Sec. II A 1)
to higher moments and distributions in stochastic reaction networks. SCH is a homeostatic condition for the distribution p(x|s) of a size-dependent stochastic process to be expressed as a mixture of Poisson random variables drawn from an underlying continuous stochastic concentration vector *κ* = (*κ*_1_, *κ*_2_,…, *κ_N_*)^*T*^ that is statistically independent of *s*:

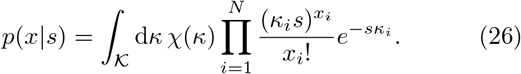

The fact that *κ* and its density *χ*(*κ*) are independent of s ensures concentration homeostasis in the stochastic sense.

### A. The effective dilution model is valid for reaction networks with stochastic concentration homeostasis

Theorem 1 (Appendix A) is a central result of our analysis and it states that if the EDM (4) satisfies SCH, i.e., Eq. (26) holds for Π_EDM_(*x|s*), then its stationary solution is also a solution of the ABM (25):

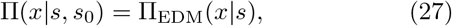

and the solution is independent of the birth size *s*_0_. Equivalently (Appendix A), we can say that the EDM/ABM satisfies SCH if the factorial-moment generating function is of the form

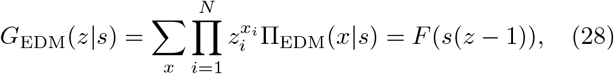

where 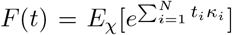, the moment-generating function of the concentration vector, is cell-size (*s*) independent. An interesting observation is that SCH implies that the mean numbers and coefficients of variation for cells of the same size *s* are given by

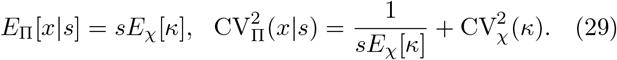

Since *κ* is independent of *s*, SCH implies homeostasis of the mean concentrations in (29) but concentration homeostasis on average does not necessarily imply SCH. The coefficients of variation coincide both for concentrations and molecule numbers and have size-dependent and size-independent components. In the following, we provide examples of reaction networks for which SCH holds for all values of the rate constants and demonstrate the validity of the EDM by comparing its distribution solutions to ABM simulations.

It can be seen from (26) and (29) that when the EDM’s stationary distribution is Poissonian with deterministic concentration vector *κ*, this distribution satisfies SCH and hence is also a solution of the ABM. More generally, SCH can be checked without solving for *G*_EDM_(*z*|*s*) (or Π_EDM_(*x*|*s*)). Assuming mass-action kinetics (6), for example, a sufficient condition for SCH is that the network consists entirely of mono-molecular reactions (see Appendix A) of the form:

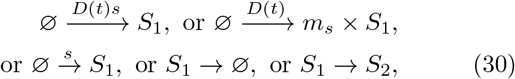

where *S*_1_ and *S*_2_ denote any pair of species that are partitioned at cell division and *D*(*t*) is a exogenous stationary stochastic process modelling a genetic state which is copied but not partitioned at cell division and does not scale with cell size. The propensities of zero-order reactions in SCH networks must either be proportional to cell size or include size-dependent random bursts *m_s_* whose burst distribution satisfies SCH itself. We illustrate the predictive power of this result by demonstrating SCH for common gene expression models involving reactions of the form (30) and show that the analytical solution of the chemical master equations agrees exactly with the agent-based models (Fig. 2a-c).

**FIG. 2.**
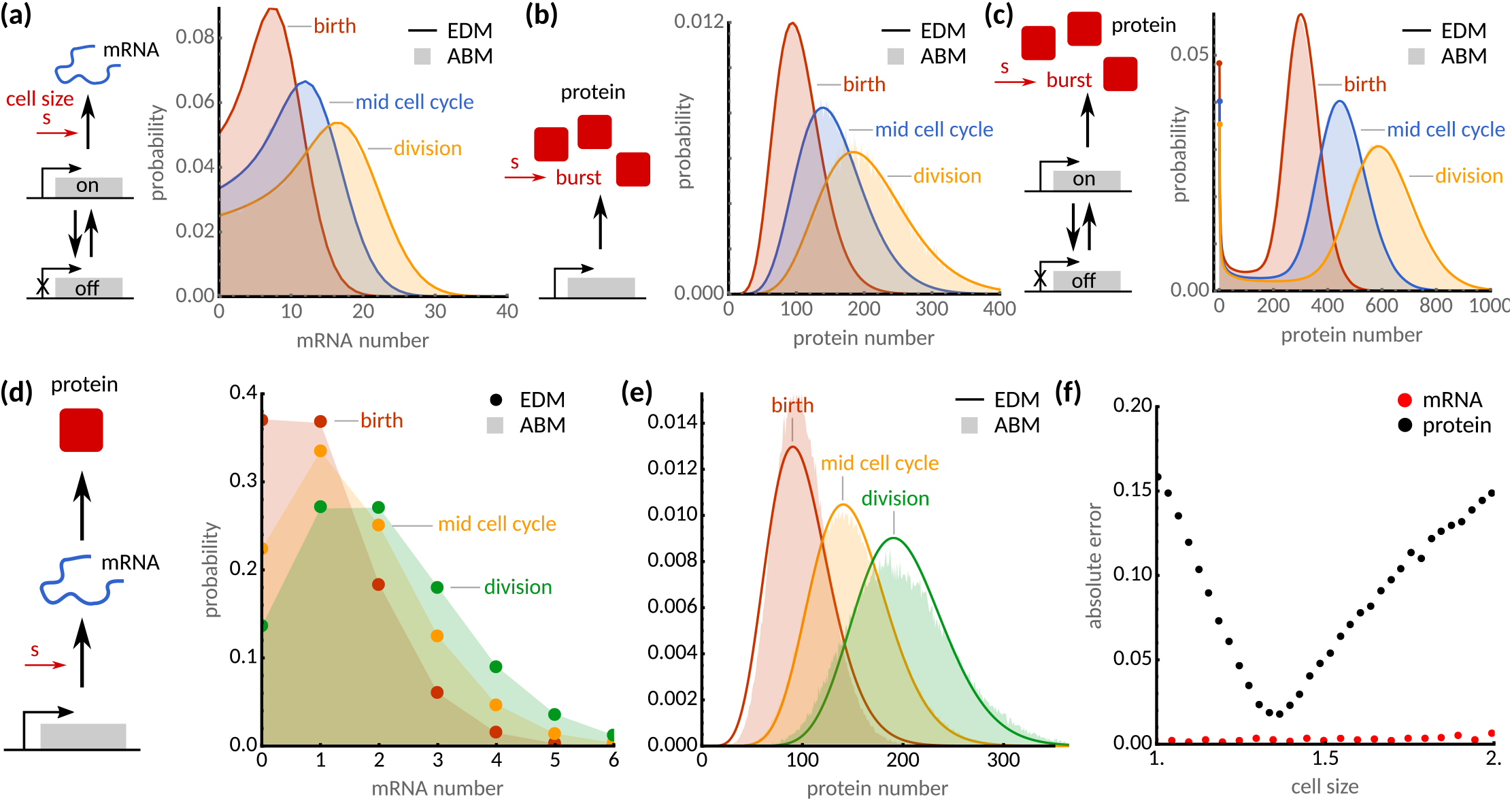
Distributions of CME and agent-based models agree for reaction networks with stochastic concentration homeostasis. **(a-c)** The EDM (solid lines, analytical solution^53,54^) agrees with agent-based simulations (shaded areas) for a range of gene expression models. Panels show (a) bursty transcription^53^ with transcription rate proportional to cell size, (b) bursty translation^54^, and (c) bursty transcription and translation^54^ with geometrically distributed bursts *m_s_* whose average is proportional to cell size *s* (see main text for details). **(d)** mRNA distributions simulated using the ABM (shaded areas) are shown for cells of sizes *s* = so (red), *s* = 1.5*s*_0_ (orange) and *s* = 2*s*_0_ (green), which agree with the effective dilution model (dots, Poisson distribution). **(e)** Simulated protein distributions (shaded areas) disagree with the effective dilution model (solid lines, solution in Ref.^55^). **(f)** Absolute error (*ℓ*_1_) of the effective dilution model as a function of cell size for mRNA (teal) and protein (red) distributions. ABM simulations were obtained using the First-Division Algorithm (Box 1) assuming an adder model (*a* = 1) and parameters *k*_0_ = 10, *k*_dm_ = 9, *k*_tl_ = 100, *α* = 1. Cell cycle noise assumes gamma distribution 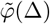 with unit mean and CV_*φ*_[Δ] = 0.1, while division noise assumes symmetric beta distribution with CV_*π*_ [*θ*] = 0.01.

mRNA expression involving a two-state promoter^53^ (Fig. 2a),

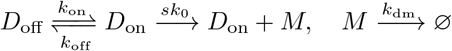

satisfies SCH for all parameter values since the network is of the form (30) whenever the transcription rate is proportional to cell size. The stochastic concentration variable is distributed as 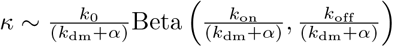.

Bursty protein expression (Fig. 2b) of a stable (nondegrading) protein arising from a two-stage model of gene expression can be modelled using stochastic bursts:

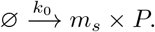

According to (30) the model satisfies SCH for all parameter values when the burst distribution obeys SCH. This is the case for geometrically distributed bursts *m_s_* whose mean is proportional to cell size, *E*[*m_s_*|*s*] = *bs*. It can be shown that the stochastic concentration variable follows κ ~ Gamma(*k*_0_/*α, b*).

Similarly, bursty protein expression from a two-state promoter arising from a three-stage gene expression model^54,56^ (Fig. 2c),

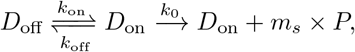

also satisfies SCH for geometrically distributed bursts (with mean *bs*) but the concentration variable is doubly stochastic^57^ 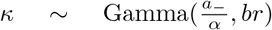, 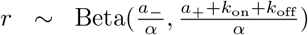 where 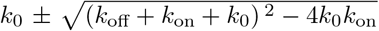. We observe excellent numerical agreement between the ABM simulations and analytical EDM solutions in all these cases validating our theoretical predictions (Fig. 2a-c).

A complex example of the reactions (30) that obeys SCH for all parameter values but yet defies analytical solution is

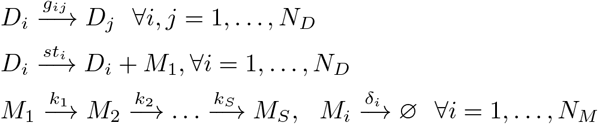

where the exogenous genetic states *D_i_* undergo switching with rates *g_ij_* but are not partitioned at cell division, transcription rates *st_i_* are assumed to be proportional to cell size *s*, and processing of transcripts *M_i_* follows a multi-step process with rates *k_i_* and degradation with rates *δ_i_*. For example, it can be checked that for *N_M_* = 1 we recover the 2^*m*^-multistate model^58^ as a special case whose EDM has a factorial-moment generating function (compare Eq. (7) in^58^ with (28)) satisfies SCH precisely when the transcription rates are proportional to cell size.

On the other hand, discrepancies between the EDM and ABM solutions will be apparent when reactions do not obey SCH. To illustrate this point, we return to the gene expression model with transcriptional size-scaling and explicit protein translation reaction (13). Note that in the EDM extra reactions are being added for the dilution of mRNAs and proteins, while for the ABM proteins are diluted through growth and divisions. Using our condition (28), it is straight-forward to verify that the Poissonian mRNA distributions of the EDM coincide exactly with the distributions of the ABM (Fig. 2d). However, this condition is not met for the protein distribution since the translation reaction is not a monomolecular reaction of the form (30). To demonstrate the breakdown of the EDM, we compare the analytical steady state distributions obtained by Bokes et al. ^55^ against ABM simulations at various cell sizes (Fig. 2e). We observe that the error of the EDM (as quantified by the *ℓ*_1_-distance of the two distributions, Fig. 2f) is pronounced both for newborn and dividing cells. The remainder of this article is dedicated to investigate the sources and consequences of these discrepancies.

### B. The EDM approximates the mean concentrations of ABMs lacking SCH

SCH provides a general criterion with which to probe the validity of the EDM probability distributions. In practice, however, approximate agreement of the first few moments, e.g., mean and variances, often suffices. Here, we establish that under the mass-action scaling assumption (6) the mean concentrations of the ABM and EDM agree approximately, and they satisfy concentration homeostasis on average. This can be seen by multiplying Eq. (25) by *x* and averaging, which yields ODEs for the mean numbers:

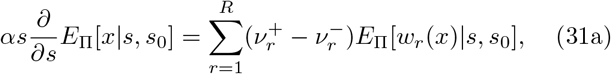

and the boundary conditions

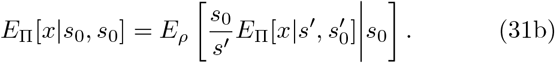

Unfortunately, Eqs. (31) are not necessarily closed since the equation for the mean may involve higher order moments when *w_r_*(*x*) depends nonlinearly on *x* and we need to resort to approximations. Analogously to the linear noise approximation, we set 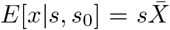 and 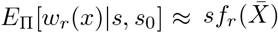 and insert the resulting expression into Eqs. (31). It follows that 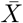 is independent of size and satisfies the rate equations (2). We conclude that, for mass-action kinetics, the EDM agrees exactly with the ABM on average for networks with linear propensities and approximately for large cell size for nonlinear reaction networks.

### C. Scaling of fluctuations with size in individual cells manifests the breakdown of the EDM lacking SCH

Next we investigate the scaling of fluctuations with cell size. Under the linear noise approximation the covariance matrix Σ(*s, s*_0_) = Cov_Π_[*x*|*s, s*_0_] evolves according to

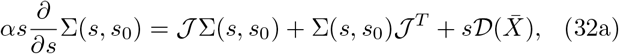

where 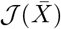 and 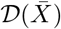 are the Jacobian and diffusion matrices defined after Eq. (11). To make analytical progress we assume for now that cell division is deterministic (CV_*φ*_[Δ] = CV_*π*_ [*θ*] = 0), which implies the following boundary condition

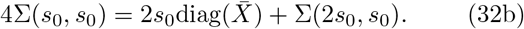

The first term is due to binomial partitioning of molecules and the second stems from gene expression noise at division. It is implicit in the deterministic division assumption (CV_*φ*_ [Δ] = CV_*π*_ [*θ*] = 0) that the birth size *s*_0_ across cells is fixed and that the size distribution in Eq. (24) reduces to

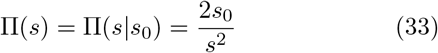

for *s*_0_ ≤ *s* ≤ 2*s*_0_ and zero otherwise, in agreement with previous results^59,60^. Similarly, the ancestral distribution (25c) reduces to 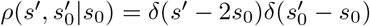.

Eqs. (32) can be solved in closed form using the eigende-composition of the Jacobian 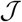. The solution to (32a) that respects the boundary condition (32b) is (Appendix D)

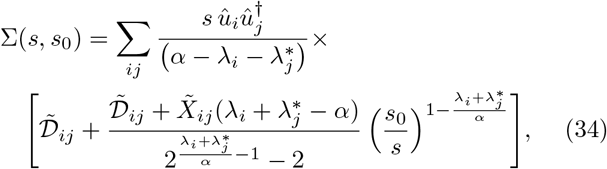

where ^†^ denotes the conjugate-transpose and we defined the matrices 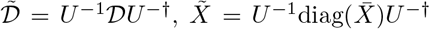 and *U* = (*û*_1_,…, *û_N_*) whose columns are the eigenvectors of 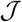 such that 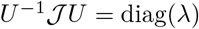.

We demonstrate the implications of this result using the gene expression example with transcriptional size-scaling and explicit translation reaction (13). The mean of mRNA numbers m and protein numbers *p* are

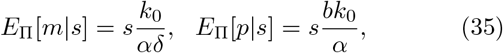

where the constants are defined in Eq. (16). These expressions hold both for the EDM and the ABM, and they exhibit concentration homeostasis on average as shown in the previous section. The exact agreement between EDM and ABM is also confirmed by ABM simulations (Fig. 3b).

**FIG. 3.**
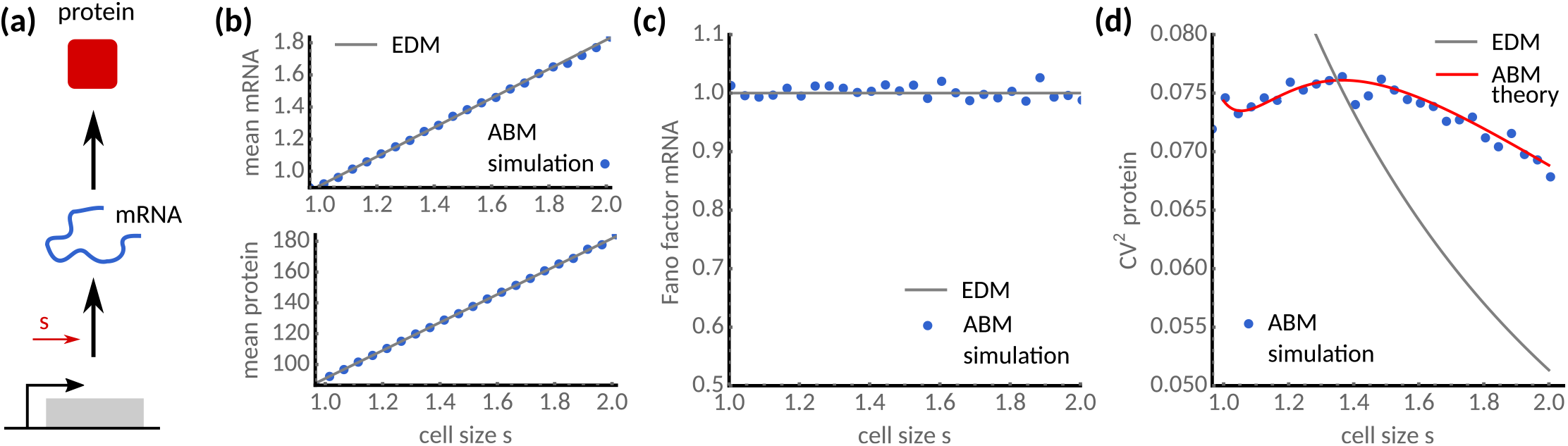
Comparing the statistics of the effective dilution and agent-based models. **(a)** Simple model of mRNA transcription and protein translation transcriptional size-scaling (13). **(b)** Mean mRNA (top) and protein levels (bottom) agree with the EDM (solid grey lines) and ABM simulations (blue dots). **(c)** mRNA statistics display unit Fano factor indicating Poisson statistics in agreement with EDM. **(d)** ABM simulations (dots, Box 1) display non-monotonic cell size scaling of protein noise, which are predicted by the agent-based theory (solid red) but not by the EDM (solid grey). Parameters are *k*_0_ = 10, *k*_dm_ = 10, *k*_tl_ = 100, *α* = 1. Cell size control parameters are as in Fig. 2.

Using Eq. (32) we find that the cell size dependent fluctuations satisfy

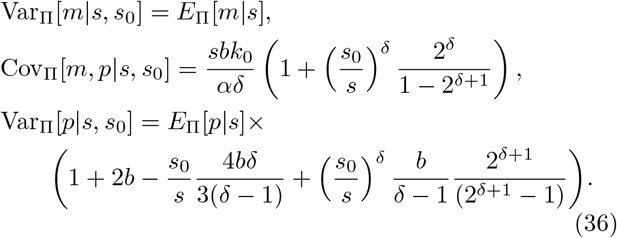

We note that the mRNA variance of the ABM agrees precisely with the EDM (Fig. 3c). The agreement is a direct consequence of SCH exhibited by the mRNA transcription and degradation reactions (cf. (30)). However, the expressions for the predicted mRNA-protein covariance and protein variance disagree with their EDM counterpart since the reactions involving the protein violate SCH. To explore this dependence, we compare the corresponding coefficients of variation of both models (Fig. 3d). The EDM overestimates cell-to-cell variation of small cells but underestimates it for large cells. Moreover, the EDM’s coefficient of variation decreases monotonically with cell size, but this is not the case for the ABM.

Strikingly, the coefficient of variation peaks as cells progress through the cell cycle (Fig. 3d, solid red line), which is in excellent agreement with the ABM simulations (blue dots) but is not seen in the EDM (solid grey). This can be seen directly from Eqs. (36) for which protein fluctuations can be approximated in the limit of fast mRNA degradation (*δ* ≫ 1) as

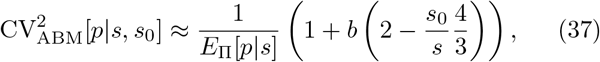

which has a maximum at a cell size of 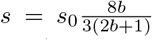 as confirmed by agent-based simulations (Fig. 3d). Depending on the burst size *b*, the peak shifts from *s* = *s*_0_ for *b* = 3/2 to *s* = 4/3*s*_0_ for *b* ≫ 1. The qualitative difference between the scalings of gene expression noise with cell size of EDMs and ABMs manifests the breakdown of the EDM, which is observed both for concentrations and molecule numbers since their coefficients of variation coincide when considering cells of the same size.

#### Effect of cell size control on gene expression dynamics

Next, we ask how fluctuations in the cell size control affect gene expression noise. It may be intuitively expected that noise in cell size control and division errors cause variable birth sizes, variable division times and hence noisy expression levels. mRNA fluctuations in the gene expression model with transcriptional size-scaling (13) obey SCH and hence are unaffected by these noise sources. The effect on protein noise remains yet to be elucidated.

To this end, we assume small birth-size variations and approximate the actual birth size with an averaged estimate *E*_Π_[*s*_0_|*s*] of the retrospective birth size for a cell of size *s*. The covariance matrix (or any other moment) can then be approximated as

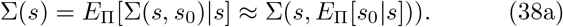

This simplification can formally be justified through a saddle-point approximation as the joint distribution Π(*s, s*_0_) has a maximum at Π(*s, E*_Π_[*s*_0_|*s*])). Generally no analytical expression of *E*_Π_[*s*_0_|*s*] can be derived from Eq. (24) in the presence of cell size control fluctuations, however, and we approximate *E*_Π_[*s*_0_|*s*] by a matched asymptotic expansion (Appendix C):

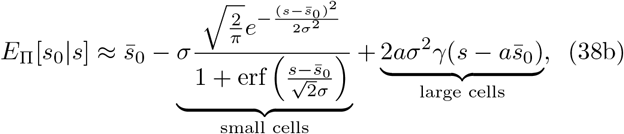

which holds for the linear cell size control model (19). The first term is the average birth size in the absence of cell size control fluctuations, the second term denotes the contributions from small cells, while the third term stems from large cells. The interpretation of this conditional expectation is that small cells were born with sizes smaller than average while larger cells were born with sizes above average depending on their size control (Fig. 6 in Appendix C). The parameters in Eq. (38b) are given by the mean birth size 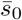 and variance *σ*^2^ in a backward lineage tracing the ancestors of a random cell in the population (see Ref.^48^ for details):

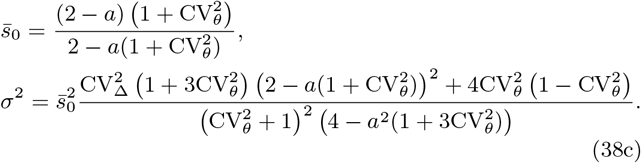

Eqs. (38) provide a closed form approximation of the cell size dependence of any given moment accurate to order *O*(*σ*^3^).

To verify the accuracy of proposed approximation, we test the theory for various strengths of size control noise and division errors (Fig. 4). We observe that increasing noise results in the monotonic decrease of gene expression noise with cell size (Fig. 4a-d) in good agreement with ABM simulations, even for large cell size fluctuations. We further ask about the effects of partitioning noise, which shows a similar dependence but agrees less well with the ABM simulations for cells smaller than the mean birth size (Fig. 4eh), presumably since the effect of large variability in birth sizes is not captured in our approximation. Nevertheless, the present approximation qualitatively captures the overall cell size dependence of the ABM simulations (Fig. 4). Our findings confirm that birth size variation contributes significantly to the cell size dependence of gene expression noise of networks lacking SCH.

**FIG. 4.**
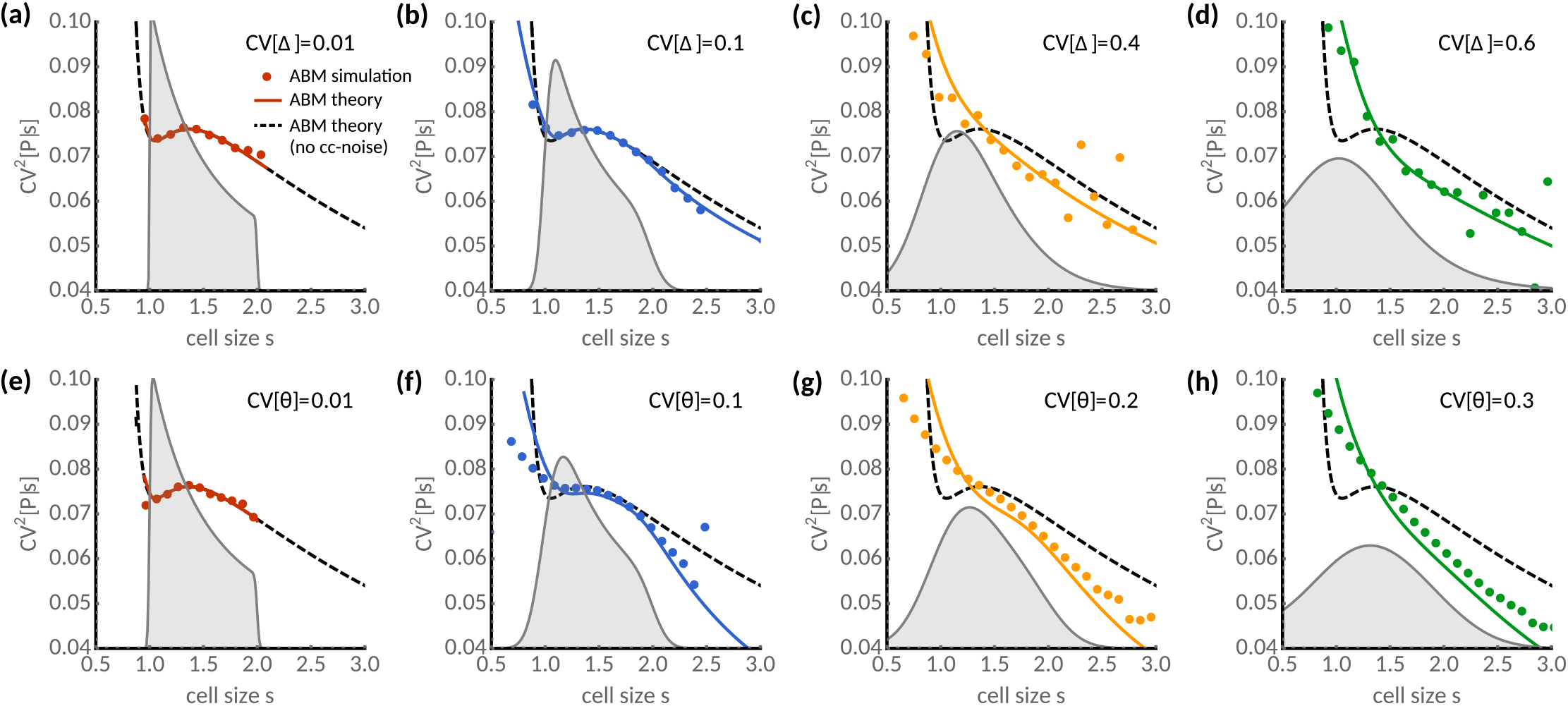
Effect of noise in cell size control and division on gene expression noise in single cells. **(a-d)** Protein noise as a function of cell size s for various noise levels in added size CV[Δ]. For comparison, the prediction without cell cycle noise (dashed black line, Eq. (36)) and the cell size distributions (shaded grey) are shown. **(e-h)** Same as (a-d) but with division noise affecting the inherited size fraction CV[*θ*]. Analytical predictions (solid lines, Eq. (38a) with (36) and (38b)) and ABM simulations (dots) using the First-Division Algorithm (Box 1) are shown. Gene expression model and all other parameters are as given in Fig. 3. Added size Δ assumes a gamma distribution with unit mean and CV[Δ] = 0.01 (e-f) while division errors *θ* followed a symmetric beta distribution with CV[*θ*] = 0.01 (a-d).

### D. ENMs provide surprisingly accurate approximations of total noise in ABMs lacking SCH

We go on to compare the ENM introduced in Sec. II with the ABM. In contrast to the EDM, the ENM predicts the total noise statistics including the variability introduced by the cell size distribution. It is clear that the ENM agrees exactly with the ABM whenever the network obeys SCH. In particular, the marginal factorial-moment generating function of the ABM’s molecule numbers 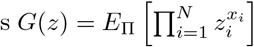 (irrespective of cell size) follows from Eq. (26) as

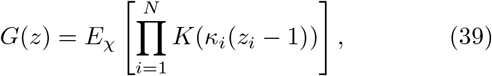

when the concentration distribution *χ* has been identified (as we did for the models in Sec. III A) and the moment-generating function *K*(*t*) = *E*_Π_[*e^ts^*] of the cell size distribution (24) is known. When SCH does not hold, the ABM statistics can in principle be obtained through integrating Eq. (34) against the size distribution Π(*s,s*_0_). Specifically, denoting molecule numbers by *x* and concentrations by 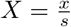, as before, we have

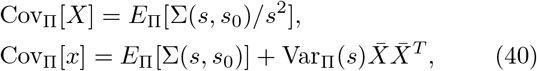

where Σ(*s, s*_0_) is the size-dependent covariance matrix discussed in Sec. III C.

To illustrate this dependence, we consider the gene expression model with transcriptional size-scaling (13) and integrate Eq. (36) numerically against the size distribution (24). We observe that the mRNA noise-mean relationship of the ABM follows exactly the ENM predictions when the mean varies through the transcription rate (Fig. 5a). This agreement is confirmed for various strengths of cell size control fluctuations and division errors, both for mRNA concentrations and numbers, which validates our theoretical predictions that the mRNA distribution satisfies SCH and hence the ENM is exact.

**FIG. 5.**
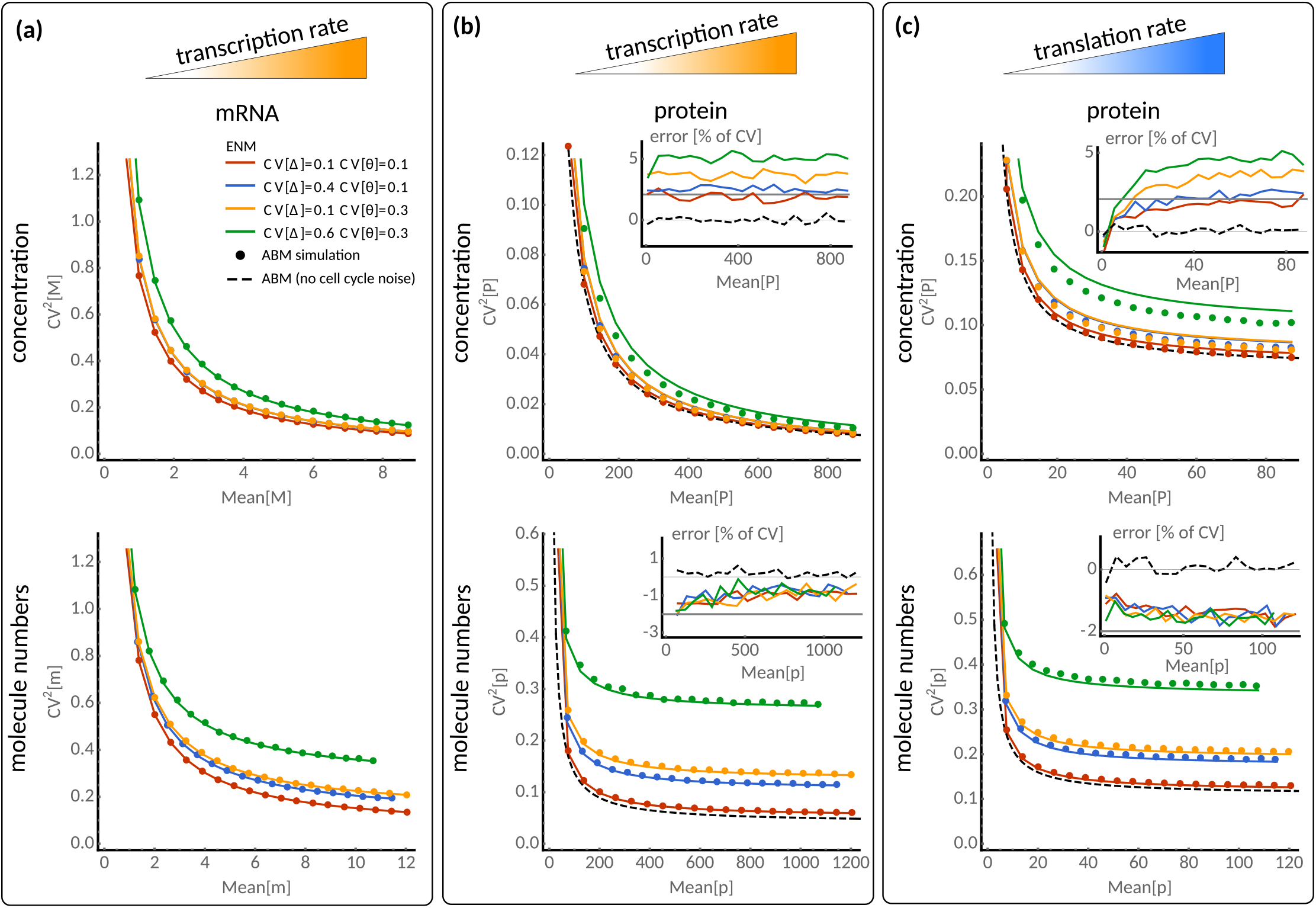
The extrinsic noise model approximates gene expression noise with size control and division errors. **(a)** Scaling of mRNA concentration noise with mean concentrations for various noise levels in added size CV[Δ] and partition noise CV[*θ*] when the transcription rate *k*_0_ is varied (top). Corresponding scaling is shown for mRNA numbers (bottom). Analytical predictions of the ENM (solid lines, Eq. (8) with (10) and (12)) and ABM simulations using the First-Division Algorithm (dots, Box 1) are shown. The inset shows the relative error in CV of the EDM compared to ABM simulations [100% × (CV_ENM_/CV_ABM_ – 1)]. **(b)** Scaling of protein noise with mean protein concentration (top) and numbers (bottom) when the translation rate *k*_tl_ is varied. **(c)** Same as (b) but varying the transcription rate *k*_tl_. For comparison, the ABM predictions without cell cycle noise are shown (dashed black lines, Eq. (41)) and the error bounds of 2% predicted by the theory (solid grey). See caption of Fig. 3 for the remaining parameters.

**FIG. 6.**
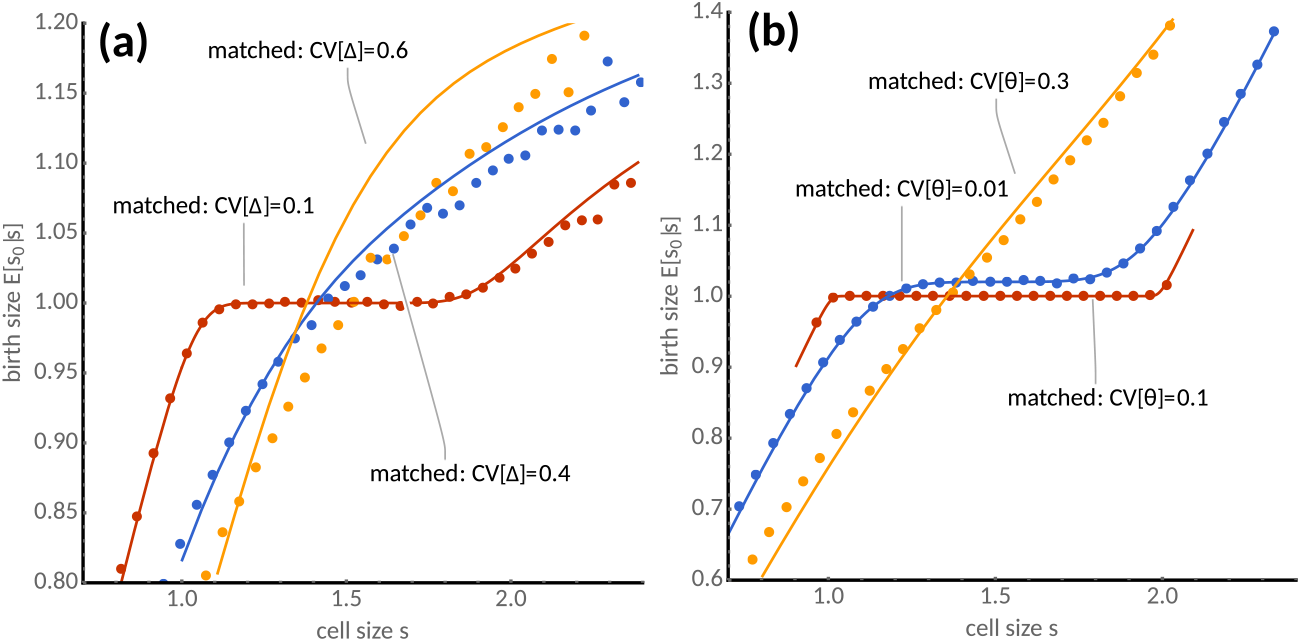
Retrospective averages of birth size. Matched asymptotic expansions of *E*_Π_[*s*_0_|*s*] (Eq. (38b), lines) for the adder size control (*a* = 1, *s*_0_ = 1) and agent-based simulations (dots) for (**a**) varying size control noise CV[Δ] and (**b**) division errors CV[*θ*].

However, the protein noise-mean relationships of the ABM and ENM differ (Fig. 5b). The discrepancy, albeit small, exists even for deterministic divisions (CV_*φ*_ [Δ] = CV_*π*_ [*θ*] = 0) for which the averages (40) over the size distribution can be carried out analytically and result in:

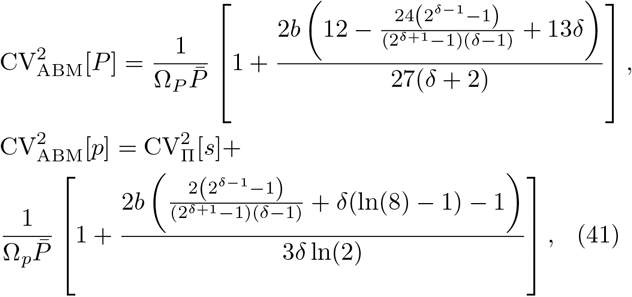

where Ω_*P*_ = *E*_Π_[*s*^-1^]^-1^ and Ω_p_ = *E*_Π_[*s*] and *δ, b* are defined as in Eq. (16). It can be verified by optimising (41) over δ that the ENM underestimates ABM noise of protein numbers by at most 2%, while it overestimates noise in protein concentrations by the same amount. The difference between the ENM and ABM predictions increases with cell size control noise for concentration measures but appears to be practically independent of cell size control noise for protein number fluctuations (Fig. 5b, insets). The protein number noise, but not concentration noise, exhibits an extrinsic noise floor (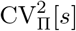 in (41)) for large mean numbers due to extrinsic cell size variability across the population and this noise floor increases with noise in size control (CV_*φ*_[Δ]) and division errors (CV_*π*_ [*θ*]) as it is also predicted by the ENM (Fig. 5b).

Similar conclusions hold for the noise-mean relationship when the mean is varied through translation rate (Fig. 5c) but there appears an additional intrinsic noise floor due to stochastic bursting (Fig. 5c and Eq. (41)) that is present both in the protein concentration and number noise. This phenomenon is in qualitative agreement with previous findings^27,35,40^ and similarly predicted by the ENM (15). Presumably, the better quantitative agreement for molecule numbers as compared to concentrations (Fig. 5b and c) is due the fact that the ENM and ABM predictions are dominated by extrinsic noise, which has the same effect in both models. Our observations suggest that, surprisingly, the ENM provides much more accurate approximations of the ABM statistics than the EDM.

## IV. DISCUSSION

We presented an agent-based framework to study gene expression noise coupled to cell size dynamics across growing and dividing cell populations. The framework consists of an exact algorithm for simulating the stochastic dynamics of dividing cells (Box 1), which generalises previous algorithms for isolated lineages^4,17,32,61–63^ towards growing cell populations, and a master equation framework (Sec. II) that exactly characterises the snapshot-distribution of gene expression and cell size across such a agent-based population.

Our theory shows that the newly defined stochastic concentration homeostasis (SCH) (cf. Sec. IIIA and Theorem 1) provides a necessary and sufficient condition for the stationary distributions of the chemical master equation (EDMs and ENMs) to agree exactly with the snapshot distributions of detailed agent-based models (ABM). A broad class of gene networks (30), involving mono-molecular reactions, multi-state promoters and bursting, satisfy SCH irrespective of network parameters when the reaction rates scale with cell size according to the law of mass-action. SCH is however not restricted to this particular class of network and can generally be checked on a case-by-case basis using the generating function equations, which can be accomplished without solving the chemical master equation analytically (see Appendix A).

SCH guarantees that gene expression distributions for cells of a given size are entirely independent of extrinsic noise sources affecting birth size such as cell size control and division noise. They thus reveal whether a network embedded in a growing cell can be insulated against such noise sources, an important feature that can guide the design of synthetic circuits.

Nevertheless, most gene regulatory networks of interest do not obey SCH. To address this issue, we developed the linear noise approximation for ABMs lacking SCH embedded in growing and dividing cells. The theory provides ODEs for the mean molecule numbers (31) and their covariances (32), which – unlike the conventional linear noise approximation describing EDMs^6,43^ – describe their evolution across sizes and features a boundary condition describing the stochastic partitioning of molecules at cell division. We showed that, when the reactions follow mass-action size-scaling, the mean concentrations of EDMs and ABMs agree, because they exhibit concentration homeostasis (Sec. IIIB). The theory further provides closed-form analytical expressions for the covariance matrix of gene expression fluctuations in the absence of SCH. We note that like the conventional linear noise approximation, the linear noise approximation for ABMs is exact for linear reaction networks but represents an approximation for nonlinear reaction networks (Sec. IIIC).

While the EDM always predicts birth-size-independent noise, the ABM’s covariance matrices generally depend on it (Sec. IIIC), both for concentrations and molecule numbers. This means that, unlike in SCH conditions, size-control noise and division errors can propagate to gene expression levels, and we unveiled quantitative and qualitative differences between EDMs and ABMs regarding the dependence of expression noise on cell size. Such differences prevail even for relatively simple gene networks involving protein expression (see (13)) for which our linear noise approximation readily provides exact expressions for mean and noise statistics.

Despite these discrepancies, we found that the ENM of these simple gene expression models provides surprisingly accurate total noise estimates (Sec. IIID). In fact, we showed analytically that the ENM (and EDM) of bursty production with translational size-scaling agrees exactly with the ABM since it obeys SCH. For transcriptional size-scaling, which implies the absence of SCH, the ENM deviates at most a few percent from ABM’s total protein noise prediction. To resolve such small differences experimentally, one would need to probe on the order of 10, 000 cells for measuring the squared coefficient of variation accurate to three leading digits (assuming sampling errors inversely proportional to cell number), which is achievable only with high-throughput techniques.

An outstanding question is whether the good agreement we observed is specific to the particular model or parameter values we have chosen or whether the ENM is more generally valid. We have made an initial step in this direction by providing closed-form expressions for the ABM’s linear noise approximation of any single-species reaction network with deterministic size-distribution (Appendix D). These results demonstrate that the ENM overestimates the ABM’s coefficient of variation of concentrations by at most 8% but underestimates it by at most 2%, and vice versa for molecule numbers, and these bounds hold independently of the choice of parameters. This suggests that ENMs could be surprisingly accurate approximation of ABMs. Other effective models of bursty protein production without any size-scaling as proposed in^64^ cannot obey SCH since they ignore cell size and generally produce larger errors than the ENM even for deterministic cell cycles.

A limitation of our study is that we assumed the validity of the linear noise approximation for the noise statistics of networks lacking SCH. Mean and covariance of the linear noise approximation are exact for linear reaction networks, as those we have studied here, but it represents an approximation for networks with nonlinear propensities valid in the limit of large molecule numbers. To improve the estimates of our theory, one could consider higher-order terms in the system size expansion^44^, resort to moment-closure approximations^65^, or to compute moment bounds^66^ for nonlinear reaction networks.

Another limitation is that we neglected growth rate variability, which is a significant source of noise at the singlecell level^47^. It would be interesting to include these features in our ABMs, compare them to the effective models, and investigate whether SCH can be generalised to this case. Previous studies^3^ have investigated the dependence of gene expression noise on growth rate dynamics in isolated lineages using small noise approximations similar to the one used here. Nevertheless, it may be expected that selection plays a pronounced role in populations where cells compete for growth unlike in isolated lineages, which in turn may lead to significant deviations of ENMs from ABMs^38,67,68^ that we have not studied here.

In summary, we proposed SCH as a general condition for exactness of EDMs. In the absence of SCH, we found that despite qualitative differences in the predictions of EDMs, ENMs closely approximate the total noise statistics of ABMs. Our results reinstate the validity of effective models as approximations of the agent-based dynamics, and thus they significantly extend the scope of state-of-the-art master equation methods to a broad range of single-cell analyses in growing cell populations.

## CODE AVAILABILITY

An implementation of the First-Division Algorithm (Box 1) in Julia is available at github.com/pthomaslab/fda.

## AUTHOR CONTRIBUTIONS

PT and VS designed the study and interpreted the results. PT developed the theory and analysed the data.

## FUNDING

This work has been supported by a UKRI Future Leaders Fellowship (MR/T018429/1) to PT and the EPSRC Centre for Mathematics of Precision Healthcare (EP/N014529/1).

## Appendix A: Stochastic concentration homeostasis and the validity of EDMs and ENMs

Appendices A and B use multi-index notation. In brief, a multi-index is a *N*-tuple *α* = (*α*_1_, *α*_2_,…, *α_N_*). One defines powers of a vector *x* via 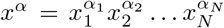, derivatives 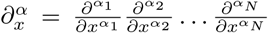, sum of components |*α*| = *α*_1_ + *α*_2_ + ⋯ + *α_N_*, and the factorials *α*! = *α*_1_! · *α*_2_! ⋯ *α_N_*! and analogously 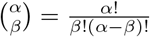.

### Definition 1

(Stochastic concentration homeostasis). *A probability mass function* Π(*x|s*) *with state space* 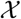 *obeys SCH if for all s* ∈ [0, ∞) *there exists a random variable κ on* 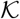 *with density χ*(*κ*) *satisfying:*

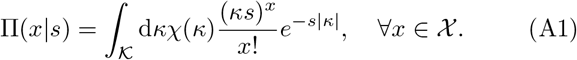

Definition 1 implies that if *κ* has a moment-generating function 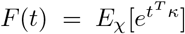 then using (A1) one finds the factorial-moment generating function

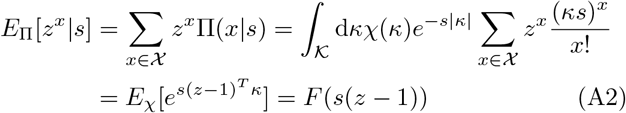

and similarly the factorial moments

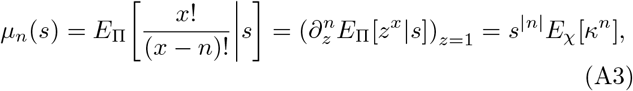

for a multi-index *n*. Since the factorial-moment generating function uniquely determines the distribution, Eqs. (A2) and (A3) may equivalently serve as definitions of SCH. Furthermore, when {*x*(*s*)}_*s*∈[0,∞)_ is interpreted as a point processes along the size coordinate *s*, SCH emphasises the fact that it is mixed Poisson process with stationary concentration vector *κ*.

### Theorem 1.

*Assume that the partitioning kernel B*(*x|x*′, *θ*) *is binomial with probability θ given by the ratio of daughter birth size and mother division size. A stationary solution of the EDM* (4), *if it exists, is also a solution of the ABM* (25):

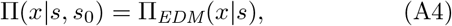

*if and only if the EDM’s solution* (4) *obeys SCH*.

The utility of the theorem is that SCH can be checked without solving the chemical master equation. We demonstrate this aspect for a general reaction network of the form (1) with mass-action propensities 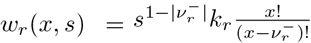, whose factorial-moment generating function (see Chapter 7 in^69^) obeys:

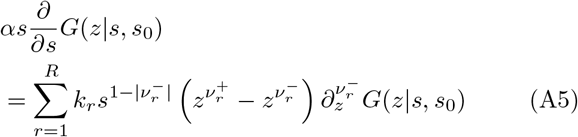

Substituting *G*(*z|s, s*_0_) = *F*(*s*(*z* – 1)) and *x* = *s*(*z* – 1) gives *αx* · ∇*F*(*x*)

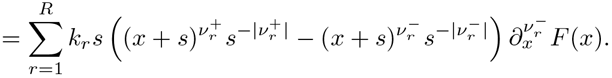

It can now be seen that the right-hand side of the above equation is independent of *s* if either (i) 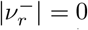 and 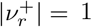, (ii) 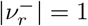 and 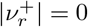, or (iii) 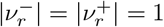. Thus the EDM and the ABM solutions coincide for mass-action reaction networks (1) when they comprise only the mono-molecular reactions given in (30).

Similarly, we check that adding bursty reactions of the form 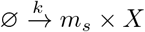 leads to a generating function equation

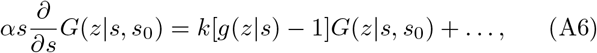

where 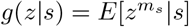 is the factorial-moment generating function of the burst distribution. Eq. (A6) transforms to *αx* · ∇*F*(*x*) = *k*[*g*(*x/s* + 1|*s*) – 1]*F*(*x*) after substituting *G*(*z|s, s*_0_) = *F*(*s*(*z* – 1)) and *x* = *s*(*z* – 1). Hence, bursty reactions obey SCH if and only if the burst distribution obeys SCH, i.e., if there exist a moment-generating function *f* satisfying *f*(*x*) = *g*(*x/s* + 1|*s*) as in (A2).

## Appendix B: Proof of Theorem 1

The proof of Theorem 1 is divided in three steps. The first step shows that a general condition (B1) guarantees snapshot distributions that are independent of birth size. We then show that (B1) satisfies the effective dilution model and reduces to SCH for binomial partitioning at cell division. The proof also clarifies that the assumption of binomial partitioning cannot be removed under biological constraints conserving the total number of molecule numbers at cell division. General conditions for the existence of the EDM’s stationary distributions have been discussed in^70^.

### Step 1: Distributions invariant of birth size

The conditional distribution Π(*x|s, s*_0_) is independent of birth size *s*_0_ if and only if

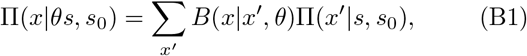

where *B*(*x|x*′, *θ*) is the partitioning kernel in Eq. (25b) that depends only the inherited size fraction *θ* = *s*_0_/*s*′. This fact can be verified using (B1) in the boundary condition (25b), which leads to

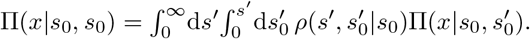

This implies that Π(*x|s, s*_0_) must be independent of birth size

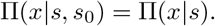

In the following, we show that under condition (B1) Π(*x|s*) coincides with the EDM solution.

### Step 2: Transformation into an effective dilution model

Let us denote the factorial-moment generating function of the partitioning kernel by *G_B_*(*z|x′, θ*) = Σ_*x*_ *z^x^B*(*x|x′, θ*) such that the invariance condition (B1) becomes

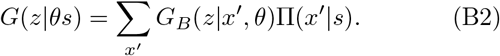

Assume that additionally

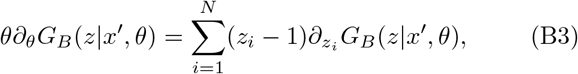

which holds for the binomial partition kernel *G_B_*(*z|x′, θ*) = (1 – *θ* + *θ_z_*)^*x*′^. Differentiating Eq. (B2) with respect to *θ* then gives

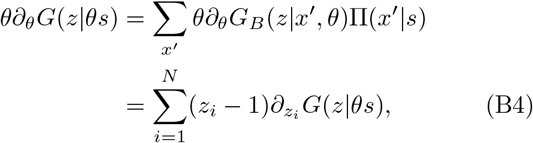

where in the last line we used assumption (B3). Changing variables (*θs* → *s*) in (B4) yields

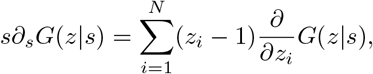

or equivalently the EDM

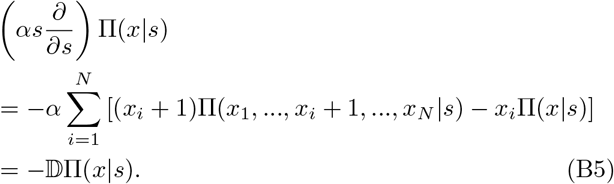

Using the above relation, we see that (25a) coincides with (4) and (A4) follows.

### Step 3: SCH and the necessity of binomial partitioning

Finally, we show that condition (B3) required for the validity of the EDM implies independent binomial partitioning of molecules. (B3) is a linear PDE that can be solved using the method of characteristics, which leads to

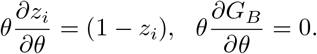

The general solution is *G_B_*(*z|x′, θ*) = *J*(1 – *θ* + *θ_z_*) where the function *J* is fixed by the condition that for *θ* = 1 all molecules are partitioned deterministically, i.e., *J*(*z*) = *z^x′^*. Hence, we obtain

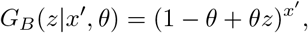

which corresponds to independent binomial partitioning of molecules in (20). It then follows that (B2) (and (B1)) are equivalent to

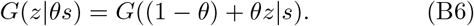

Finally, we show that (B6) is equivalent to *G*(*z|s*) = *F*(*s*(*z* – 1)) for binomial partitioning. Expanding (B6) around *z* = 1 and identifying the series coefficients with the factorial moments *μ_n_*(*s*) in (A3), we find that the factorial moments are homogeneous functions of order |*n*| = Σ_*i*_ *n_i_*: *μ_n_*(*θ_s_*) = *θ*^|*n*|^*μ_n_*(*s*). Then by Euler’s homogeneous function theorem, it follows that the factorial moments with index *n*, satisfy 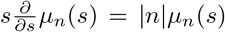 and hence *μ_n_*(*s*) = *s*^|*n*|^*μ_n_* (1). This implies that the factorial-moment generating function is

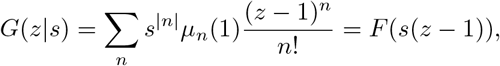

with *F*(*x*) = Σ_*n*_ *x^n^μ_n_*(1)/*n*!. It remains to be shown that *F* is indeed a moment-generating function. To this end, we note that 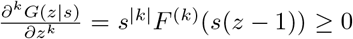 for *z* ∈ (1, −∞) and hence *F*(−*x*) is a completely monotone function on *x* ∈ (0, ∞), which implies that there exists a distribution *χ* for which *F*(−*x*) = *E_χ_*[*e^−sx^*] is a Laplace transform, which concludes the proof of Theorem 1.

## Appendix C: Approximation of birth size moments

We here derive an analytical approximation (38b) for the conditional birth size moments. We start by rewriting *E*_Π_[*s*_0_|*s*] in terms of the backward lineage distribution *ψ*_bw_ using Eq. (24):

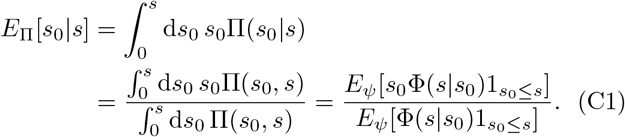

We now apply matched asymptotic expansion to this expression.

### 1. Large cell asymptotics

For large cells *s* ≫ *s*_0_, we can extend the range of integration in Eq. (C1) and compute the expectation value as follows

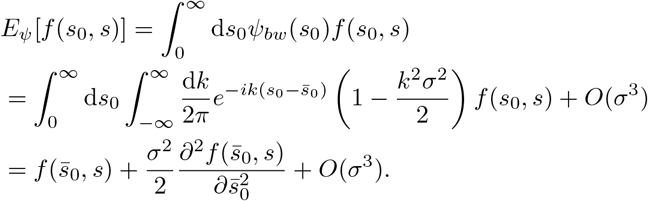

Using *f*(*s*_0_, *s*) = *s*_0_Φ(*s|s*_0_) and *f*(*s*_0_, *s*) = Φ(*s|s*_0_) in Eq. (C1), the conditional moments of birth size can be approximated by

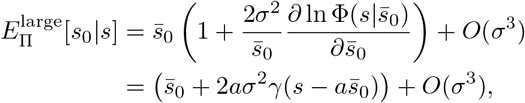

where the last equality follows from *γ*(*s, s*_0_) = *γ*(*s – as*_0_) for the linear cell size control model (19), and 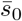 and *σ* are the mean and standard deviation of the backward lineage distribution *ψ_bw_* given by Eqs. (38c).

### 2. Small cell asymptotics

Next we consider small cells by noting that Φ(*s|s*_0_) is practically constant when *s* ≈ *s*_0_, the integral in (C1) can be approximated by

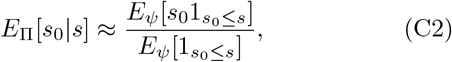

Assuming that *ψ*, is approximately Gaussian with mean 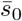 and variance *σ*^2^, we find that near 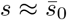, we have

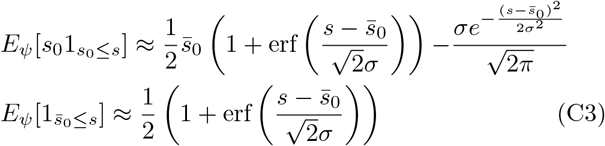

and hence

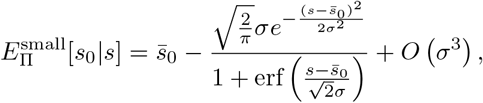

which is accurate to order *σ*^3^.

### 3. Global asymptotics

The two asymptotic solutions can be matched at the boundary layer. Since

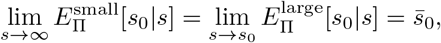

the uniformly valid matched asymptotic expansion is

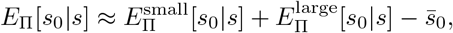

which gives Eq. (38b).

## Appendix D: Analytical solutions and error bounds using the linear noise approximation for deterministic cell division

We begin by outlining the solution of (32). Defining 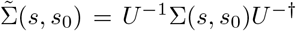, Eq. (32) of the main text becomes

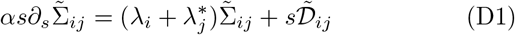

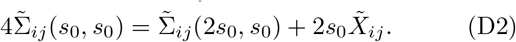

Eq. (D1) has the solution

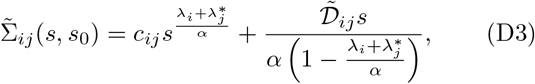

where the constants *c_ij_* are fixed using the boundary condition (D2) which gives the solution of the EDM, Eq. (34) of the main text.

Using Eq. (40) and averaging (D3) over the deterministic size distribution (33), we find the covariance matrix of concentrations *X*,

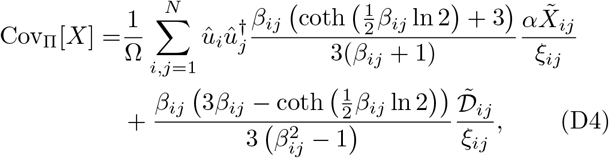

where 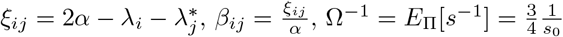, and *û_i_* are the eigenvectors of the Jacobian 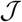 introduced before Eq. (34). Similarly, considering molecule numbers *x*, we have 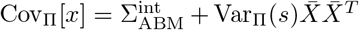 where the intrinsic noise contribution is given by

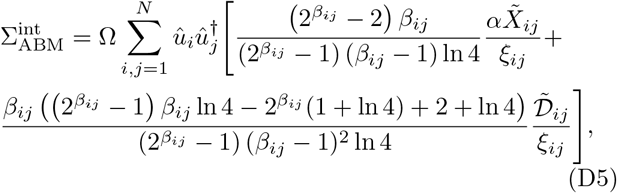

with Ω = *E*_Π_[*s*] = *s*_0_ ln 4.

The expressions greatly simplify for a single species since *û* = *û_i_*, *β* = *β_ij_* and *ξ* = *ξ_ij_*. We note that in this case (D4) increases monotonically with *β* while (D5) decreases monotonically with *β*. Using the limits *β* → 0 and *β* → ∞, we find that the ABM’s coefficients of variation can be bounded by the EDM’s coefficients:

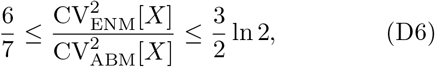

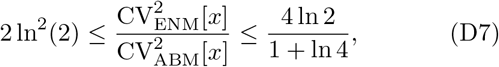

where we have used the fact that 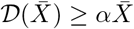 and the ENM solution of (9) is 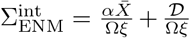. The result implies that the ENM overestimates the ABM’s coefficient of variation of concentrations by at most 8% but underestimates it by at most 2%, and vice versa for molecule numbers.

## Notes

### Competing Interest Statement

The authors have declared no competing interest.

